# Microfluidic interfaces for chronic bidirectional access to the brain

**DOI:** 10.1101/2023.09.22.558790

**Authors:** Simone Marcigaglia, Robin De Plus, Charysse Vandendriessche, Marie-Lynn Cuypers, Jordi Cools, Luis Diego Hoffman, Roosmarijn Vandenbroucke, Maarten Dewilde, Sebastian Haesler

**Affiliations:** Neuroelectronics Research Flanders (NERF), Leuven, 3000, Belgium; Department of Neurosciences, KU Leuven, Leuven, 3000, Belgium; VIB-UGent Centre for Inflammation Research, Gent, 9052, Belgium; Thermofisher Scientific (AIG/MSD), Dilbeek, 1702, Belgium; SWave Photonics, Leuven, 3000, Belgium; Laboratory for Therapeutic and Diagnostic Antibodies, Department of Pharmaceutical and Pharmacological Sciences, KU Leuven, Leuven, 3000, Belgium; PharmAbs-The KU Leuven Antibody Center, KU Leuven, Leuven, 3000, Belgium

## Abstract

Here, we used micron-scale 3D printing to develop microfluidic interfaces which provide chronic fluidic access to the brain of preclinical research models. In mice, we show the delivery interface enables faster, more precise and physiologically less disruptive fluid injection. Moreover, we demonstrate the blood brain barrier (BBB) is intact after chronic implantation of the sampling interface and establish frequent, longitudinal sampling of CSF and biomarkers from the ventricle over long time periods of up to 200 days.

## Introduction

Despite their wide importance for neuroscience and neuropharmacology, techniques which provide fluidic access to the brain are still underexplored and underdeveloped. Direct intracranial injection of chemicals or viruses, also known as convection-enhanced delivery (CED), is plagued by the tendency of the infusate to take the path of least resistance and flow back along the insertion track^1^. This phenomenon, known as backflow, aggravates with increasing flow rates^2^. Catheter miniaturization^2,3^, flexibility^4^ and porous tips^5^ have separately demonstrated some progress but have not yet been combined in one platform. Accordingly, intracranial injections today still suffer from limited predictability of fluid spread, uneven fluid distribution and long injection times.

Access in the other direction, often referred to as liquid biopsy sampling, is prohibited by the blood-brain barrier (BBB) and the blood-cerebrospinal fluid barrier (BCSFB), respectively. The gold-standard for collecting cerebrospinal fluid (CSF) in preclinical research is cisterna magna puncturing, an invasive procedure that grants ventricular access only once every two/three months^6,7^ and has a high risk for blood contamination, which yields non-representative samples. Chronic cannulation approaches are technically demanding, do not provide direct access to the ventricle and their long-term stability has not been well characterized^8–11^. Microdialysis ^12,13^ and cerebral open flow microperfusion (cFOM)^14^ on the other hand, cause significant tissue damage^15^ due to their relatively large size (∼0.5mm in diameter) and provide highly diluted samples which necessitate advanced downstream analytical methods. Microdialysis is further restricted to sampling smaller molecules below a molecular weight cut-off.

To address the aforementioned limitations and enable chronic fluid delivery (Fig. 1) and ventricular CSF sampling (Fig. 2), we describe here a novel microcatheter platform composed of specifically designed micronozzles attached to medical grade tubing. The micronozzles are fabricated using two-photon polymerization, a direct laser writing technique that allows the integration of complex, micron-scale 3D features with high precision. This combination of miniaturization and complexity cannot be achieved with conventional monolithic stacking techniques used previously for catheter microfabrication^16^.

**Figure 1.**
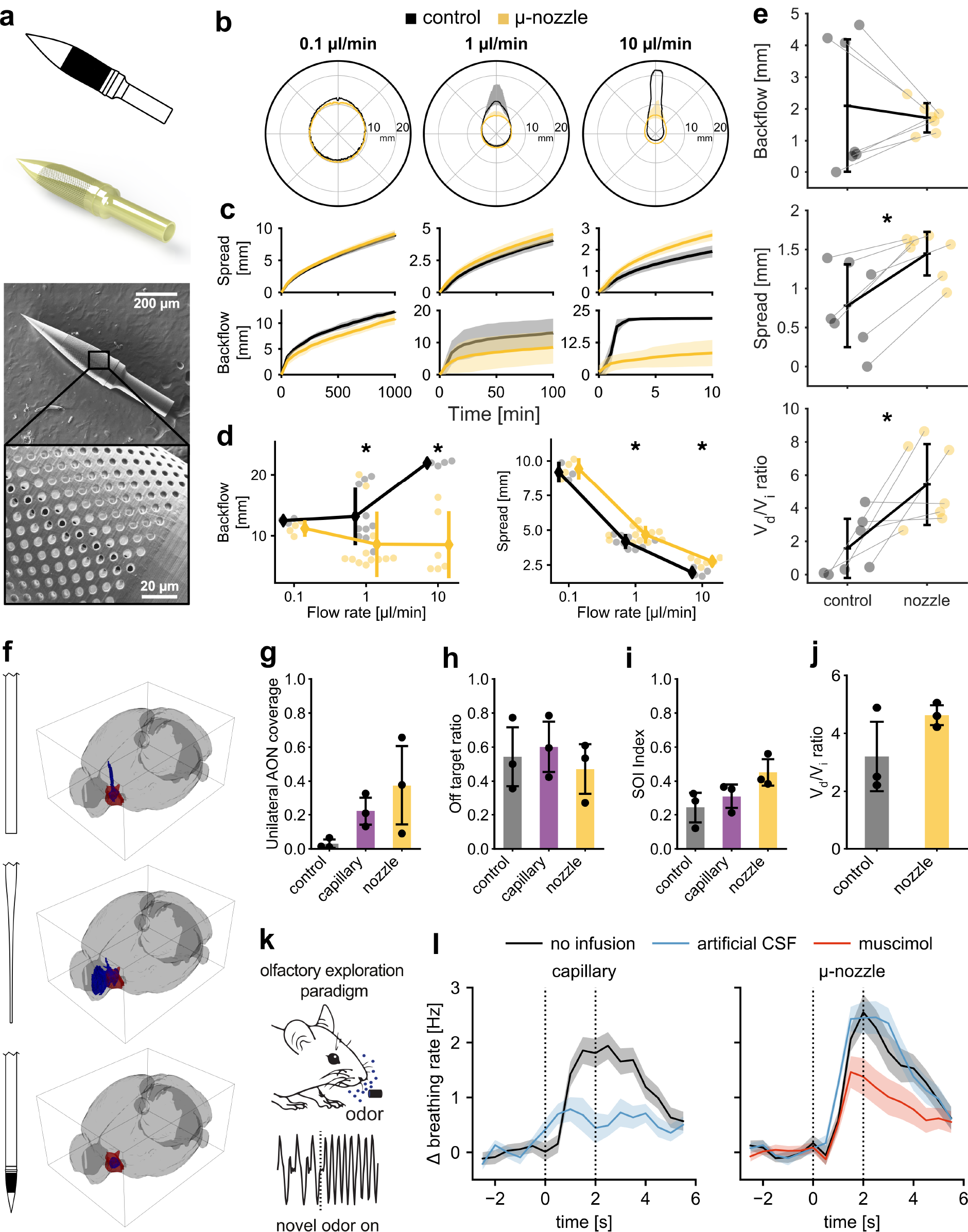
Performance of the catheter for convection-enhanced brain infusions. (a) CAD overview, 3D render of the design of the micronozzle and SEM images including a zoomed-in view of the porous region. (b) Summary of infusions with 100µl Trypan blue dye in agarose gel. Infusions are grouped by column according to flow rate. Average final shape (median±IQR) of the distribution cloud centred at the tip. (c) Average distance reached by the cloud in the lateral and downwards directions. (d) Distance travelled by the dye from the tip of the catheter along the insertion track. (e) Backflow, spread and distribution volume (VD) to infusion volume (VI) ratio at the end of the low-volume infusions in agarose gel. Infusions performed with the same protocol (delivery volume and flow rate) are joined by a line. (f) Example reconstructions of the delivery clouds (blue) for infusions performed with the control FEC [top], a glass capillary [middle] and the micronozzle [bottom]. The target structure (AON) is shown in red. Quantification of *in vivo* infusions in terms of their (g) overlap with the target structure, (h) off-target ratio and (i) overall success of infusion (SOI). (j) Ratio of distribution volume (VD) to infusion volume (VI) for dye deliveries targeted to the striatum. (k) Illustration of olfactory exploration paradigm and representative breathing response to a novel odorant. (l) Breathing rate modulation (mean±sem) after no infusion, aCSF or muscimol infusion with either acute capillary injections (left) or chronic micronozzle implantations.

**Figure 2.**
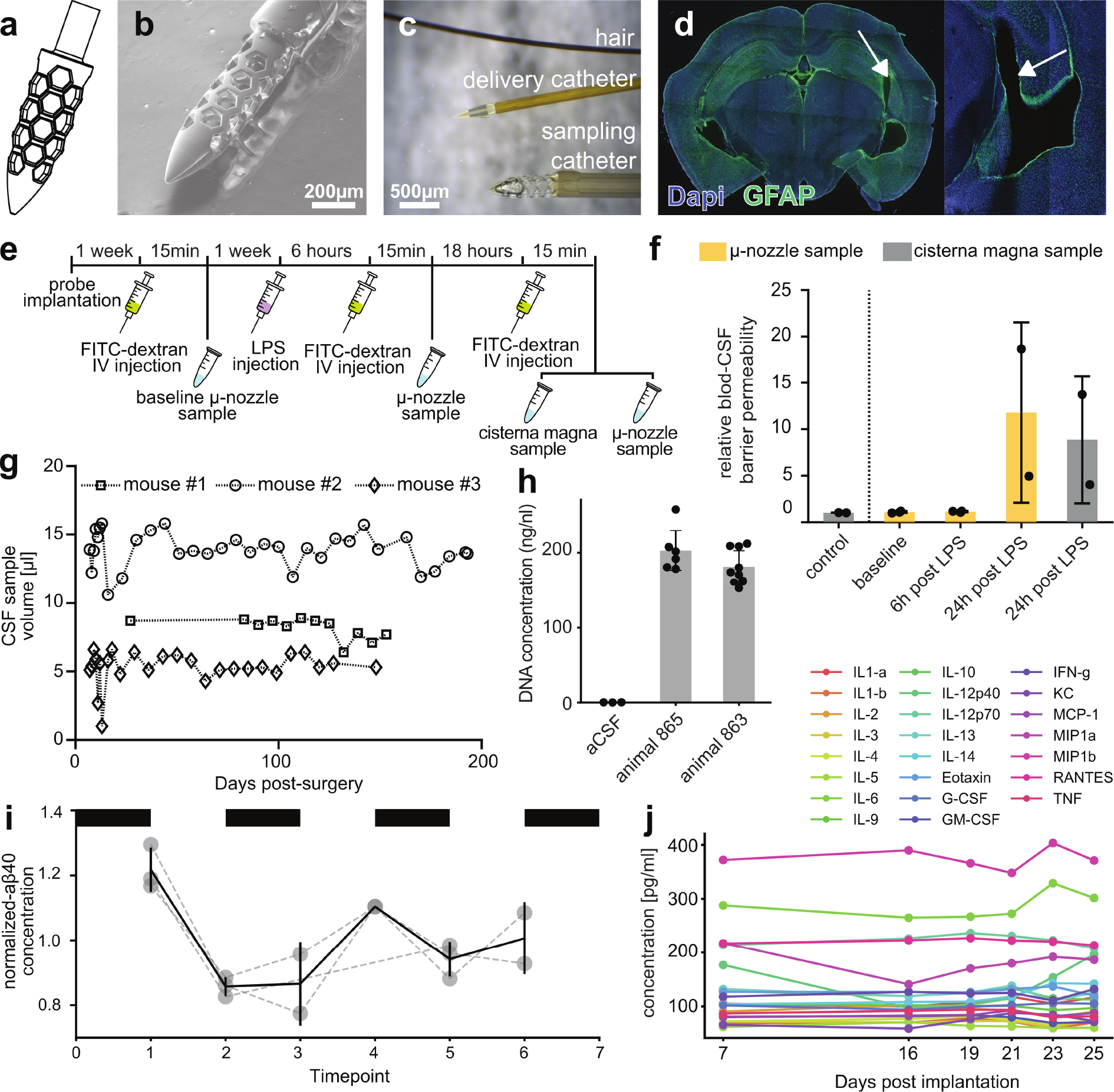
Long-term CSF sampling from the ventricle. (a) CAD design of the sampling micronozzle. (b) SEM image of the sampling unit assembled on custom 300µm-OD PEEK tubing. (c) Size comparison between a hair, the delivery and the sampling catheter, respectively. (d) Immunohistochemical staining for glial fibrillary acidic protein (GFAP, green) and nuclear DNA (blue) after chronic implantation. Arrows indicate the trajectory of entry into the ventricle. (e) Timeline of the experimental protocol used to assess the effect of sampling on BBB permeability. (f) Relative fluorescence in CSF samples. (g) Quantification of sampled volumes from three animals (mean±std). (h) Microvolume spectrophotometer quantification of nucleic acid content in 15 liquid biopsies from two animals and 3 negative controls. (i) Fluctuations in CSF levels of Aß40 at different times of the day and over days. Consecutive samples from the same animals are taken 36 hours apart. Black bars indicate night cycle (mean±std). (j) Longitudinal cytokine quantification in mouse CSF (mean±std).

The delivery micronozzle (Fig. 1a) was designed to minimize fluidic resistance between catheter outlets and brain tissue. It has an overall outer diameter of 140 µm and features 1872 pores with a diameter of 4 µm each, which constitute a six-fold increase in outflow area compared to an equally sized flat-end catheter (FEC). The sampling micronozzle (Fig. 2a-c) was designed for ventricular implantation and longitudinal CSF sampling, driven by capillary-forces and not requiring any active pumping. We chose a slightly larger diameter of 300µm and cage-like structure to facilitate capillary extraction and blockage-free insertion. Catheter tubing was chosen to match the outer diameter of the delivery and sampling micronozzles, respectively.

## Results and Discussions

We first validated the delivery micronozzle by dye infusions (100µl) in an agarose gel brain surrogate (Supplementary Fig. 1) at flow rates spanning three orders of magnitude (0.1 µl/min, 1 µl/min, 10 µl/min). We quantified spread and backflow after infusion and compared them to equal-diameter FECs. Convective processes are dominant only at the higher flow rates above 0.1 µl/min (Fig. 1b and Supplementary Fig. 2). At these higher flow rates of 1 and 10µl/min we observed a significant difference for both spread (Fig. 1c; 1µl/min: *P*<0.03, 10µl/min: *P*<0.02, Mann-Whitney U-test) and backflow (Fig. 1d; 1µl/min: *P*<0.01, 10µl/min: *P*<0.01, Mann-Whitney U-test). A second series of experiments was performed with delivery volumes (0.5-2.5 µl) and flow rates (0.1-1 µl/min) comparable to infusions in the rodent brain (Supplementary Table 1). At this smaller scale, control FEC experiments followed a bimodal distribution dictated by whether the threshold pressure for infusion was reached. If it was not, no dye exited the catheter, and no cloud was formed. This indicates that tissue coring and catheter clogging upon insertion are huge failure factors for infusions with FECs. The micronozzle yielded more predictable infusions (Fig. 1e and Supplementary Fig. 3), which showed lower variance in backflow, overall greater spread (*P*<0.01, Wilcoxon signed rank test) and larger volume of distribution (*P*<0.03, Wilcoxon signed rank test). Collectively these experiments demonstrate that the addition of the micronozzle improves the localization of the infusate *in vitro*.

To validate the performance of the delivery micronozzle *in vivo*, we performed dye infusions in the mouse brain. We compared three different delivery systems: the micronozzle, the control FEC described above and a pulled glass capillary, typically used to inject agents to the rodent brain (Fig. 1f). In a first set of experiments (n=3 mice), dye was infused in the left Anterior Olfactory Nucleus (AON, Supplementary Figure 4). For each infusion, we computed the intersection of the three-dimensional distribution clouds with the target region. The micronozzle increased target coverage, decreased off-target coverage and improved a combined success of infusion index (SOI, Fig. 1g-i). This index reflects the relationship between the distribution volume and the target region ranging from no overlap (SOI=0) to full overlap (SOI=1). The dynamics of inline pressure during infusion reflect the improved flow of fluid through the micronozzle compared to the FEC (Supplementary Fig. 5). Six additional infusions were performed in the striatum (Supplementary Fig. 6), a larger area of brain tissue than the AON. Delivery through the micronozzle also resulted in more predictable distribution cloud volumes (Fig. 1j).

The activity of neurons in the brain is highly sensitive to mechanical stress as demonstrated by the transient silencing of neural activity following the insertion of a neural probe^17^. We investigated the acute impact of fluid injection on neuronal physiology and behavior using an olfactory exploration paradigm in mice^18^ (Fig. 1k). When mice are exposed to novel odorants, they rapidly increase their breathing rate, also referred to as exploratory sniffing (Fig. 1l). Injection of artificial cerebrospinal fluid (aCSF), a pharmacologically inactive agent, into the AON through a conventional glass capillary prevented exploratory sniffing in response to novel odorants (n=6 mice). However, when the same volume aCSF was delivered through a chronically implanted micronozzle (n=6 mice), mice showed the spontaneous behavioral response also observed when no injection occurred. Injection of the GABAA receptor agonist muscimol through the implanted micronozzle on a consecutive day further revealed that novel odor-induced exploratory sniffing can be disrupted by the inhibition of neural activity in the AON (Fig. 1l). These experiments highlight an often-overlooked aspect of fluid injection and demonstrate that the micronozzle enables physiologically more favorable injections.

Next, we validated the sampling micronozzle (Fig. 2a-c) *in vivo* with a chronic implantation targeting the ventricle. One week after implantation, we confirmed that the BBB was not compromised using fluorescently labeled low-molecular weight marker fluorescein isothiocyanate-dextran (FITC-dextran) injected intravenously (Fig. 2e-f). Ventricular CSF samples obtained through the sampling micronozzle did not show any fluorescence, indicating no FITC-dextran crossed the BBB. However, when we chemically disrupted the BBB by injection of lipopolysaccharide (LPS), FITC-dextran was detected. Samples collected for comparison from the cisterna magna confirmed those results. Post-mortem immunohistochemical analysis of implanted brains further showed no evidence of local tissue inflammation (Fig. 2d). Temporarily sealing the catheter after each sample collection, we repeatedly sampled over long periods for up to 200 days (Fig. 2g), demonstrating the long-term stability and safety of the implant. We used the ability to repeatedly sample CSF from the ventricle to demonstrate longitudinal monitoring of CSF biomarkers including nucleic acids, 40 residue amyloid-beta (Aß40) and cytokines (Fig. 2h-j). Collectively, these experiments demonstrate that chronic implantation of the micronozzle allows unbiased recurrent sampling of CSF biomarkers over long time periods.

In conclusion, we developed a microcatheter platform for fluid injection and sampling which will be invaluable for studies of brain biology and disease progression in models of human disorders in rodents and other preclinical models. Since our methodical approach is based on advanced 3D printing, it is highly versatile and can be adapted to include additional functionalities in the future^19^.

## Materials and Methods

### Micronozzle design

The outer diameter of the delivery micronozzle is 140 µm with a total length of 900 µm. The porous delivery area is around 300 µm long and features 1872 pores of nominal size 5 µm and pitch 6 µm. The total outflow area is 3.68×10^4^ µm^2^, which represents an increase by a factor of 8.32 compared to an end-port catheter (FEC) with the same inner diameter (75 µm). The actual pore size (as measure by SEM) is closer to 4.2 µm, which ultimately reduces the area ratio to 5.87. The micronozzle wall thickness decreases in the porous region along the direction of flow to counteract the problem of preferential flow in proximal holes reported in previous literature.

The outer diameter of the sampling micronozzle is 300 µm with a total length of 1.2 mm. The catheter tip for CSF sampling was designed to improve insertion properties and long-term viability. The exclusive presence of side openings helps to prevent clogging of the catheter upon insertion and, at the same time, ensures that enough surface area is used for liquid biopsy uptake.

### Microfabrication

The catheter tips used for fluid delivery and CFS sampling were both fabricated with a Photonic Professional GT (Nanoscribe) in a biocompatible and non-cytotoxic proprietary resin (IP-Dip - Nanoscribe). The delivery micronozzle was assembled on extruded polyimide tubing (ID: 0.0039±0.0002”, Wall: 0.0008±-0.00025” - Microlumen) and secured by means of a UV- curable adhesive (NOA68 - Norland products). The same adhesive was used to secure the sampling catheter tip to suitably sized PEEK tubing (Extruded Sub-Lite-Wall®, ID: 0.007±0.001”, OD: 0.012±0.001” - ZEUS).

### Agarose gel dye infusions

Agarose gel was prepared at 0.4wt\% by dissolving Agarose powder (Sigma-Aldrich) in 1x TAE buffer (Sigma-Aldrich). Before solidification, the gel was poured in custom made 5×5×5cm acrylic moulds. The liquid was then allowed to cool down to room temperature while covered and it was used within 9-24 hours. A trypan blue solution was prepared by dissolving Trypan blue powder (Sigma-Aldrich) in dH2O at a concentration of 0.04wt%.

Infusions in agarose gel were performed with a dual-drive syringe pump (33DDS - Harvard Apparatus). A series of fittings and adapters (Nanotight - IDEX \& Captite - Labsmith) were used to connect a Luer-lock gas-tight syringe (1700 Series - Hamilton) first to 360µm PEEK tubing (Labsmith) and then to ∼140µm polyimide tubing (Microlumen). A tee interconnect was placed inline to allow the connection of a pressure sensor (uPS series - Labsmith). FEP tubing sleeves (IDEX) were used for all non-standard tubing sizes. Custom Matlab and Python interfaces were created to interface with both the syringe pump (for two-way control and readout) and the pressure sensor (for readout).

The progression of the infusion was recorded with a 5MP RGB camera (mvBlueFOX3 - Matrix Vision) but the optics varied between high and low-volume infusion. For high-volume (100µl) infusions, a bi-telecentric lens (TC23064 - Opto Engineering) was used to frame the entirety of the mould while keeping distortion to a minimum. In the case of low-volume (0.5-2.5µl) infusions, a microscope (NexiusZoom EVO - Euromex) was employed to achieve higher magnification. In both cases, a dichroic mirror flash unit (RT-CAS2-00-100-X-W-24V-FL - Opto Engineering) was used to ensure synchronised and even illumination of the mould. Due to the short focal length of the objective, this unit was placed next to the mould instead of in between the optics and the mould for the low-volume experiments.

The catheters were lowered into the agarose gel with a linear microactuator (Model 2660, David Kopf Instruments) at a constant speed of 4mm/s until they appeared in frame approximately at the centre of the mould. The position of the tip was recorded with the previously mentioned script as a set of coordinates relative to the captured images. An interval of 10 minutes was then allowed between the insertion of the catheter and the beginning of the infusion. Similarly, a 10-minute interval was respected between the end of infusion and the retraction of the catheter.

The sequence of images recorded for each infusion was then binarized to extract the progression of the distribution cloud with time. The backflow distance was calculated as the distance between the tip and the maximum reach of the dye along the catheter. The spread was calculated as the average distance reached by the dye in the bottom portion of the images (i.e., further in the gel than the insertion) away from the tip.

### Surgical procedures

#### Surgical preparation

Mice were anesthetized with a ketamine and medetomidine mixture (60mg/Kg and 0.5mg/Kg of body weight intraperitoneally). After successful induction, the eyes were moistened with an ophthalmic ointment (DuraTears, Alcon) to avoid dehydration and the head was shaved. Mice were then placed on a heating pad at 37°C (Harvard apparatus) to avoid hypothermia. With the mice head-fixed in a stereotactic frame (Narishige Instruments), iso-betadine was applied on the naked skin and a local anaesthetic, Xylocaine (50 µl, 20mg/ml) was injected subcutaneously under the scalp. A longitudinal incision was made along the top of the head and a window was cut to expose the underlying skull.

In the case of chronic implantations, two more steps followed. Firstly, polyacrylamide (Vetbond) was applied at the edges of the skin to limit irritation and to prevent infections. Secondly, the skull was roughened by means of a dental drill or a scalpel to improve the final adhesion of dental cement (Super-Bond C&B).

#### Creation of a surgical window for access to the AON

All mice were surgically implanted with a stainless-steel head plate to permit head-restraining during behavioural experiments and a bilateral craniotomy was performed to be able to inject/infuse muscimol into the AON. The coordinates of the craniotomy were marked (2.70mm ventrally from bregma and 1.0mm laterally from sagittal suture). Dental cement (Super-Bond C&B) was used to secure a custom-made stainless steel head plate to the skull. Cement was applied on the whole open skull, except from the region between bregma and the frontal suture. Once the cement was dry, a bilateral craniotomy was performed with the dental drill, by making concentric circles around the marked coordinates and by thinning the edges out.

Sterile saline was used to avoid drying out of the cortex during the surgery. Afterward, the craniotomy was covered with artificial dura (Dura-gel Cambridge Biotech) and a silicone cap (Quick Cast, World Precision Instruments). The artificial dura was used to avoid drying out of the cortex after surgery.

#### Micronozzle chronic implantations

After performing a craniotomy at the same coordinates as described above, a microfabricated catheter (previously sterilised with 70% ethanol and subsequently flushed with artificial cerebrospinal fluid) was inserted at a depth of 2.6mm. The insertion was performed with a manual hydraulic actuator (Narishige International). Once the catheter was lowered to the AON, a first layer of 3-component dental cement (Super Bond C&B) was applied to secure the catheter to the skull and to cover the remaining space of the burr hole. Once the cement had fully cured (∼10 minutes) and the catheter was firmly secured to the skull, it was removed from the stereotactic frame. A set of fluidic fittings (compression sleeve – PostNova, CapTite One-Piece Fitting - LabSmith, CapTite Interconnect I - LabSmith) was mounted onto the tubing to secure a strong leak-free fluidic connection. More dental cement was used to secure these fittings to the skull and to cover the entirety of the incision to prevent any entry point for bacteria. Once all the previously applied cement had cured (∼10 minutes) the fluidic port was temporarily sealed with a screw-on cap (CapTite one-piece plug - LabSmith). The absence of air bubbles within the fluidic port is ensured by flooding the adapters with aCSF prior to the fitting and sealing.

#### Implantations of cerebrospinal fluid sampling probes

Wild-type mice were prepared for surgery as explained above. A craniotomy was performed stereotactically to gain access to the lateral ventricle (at either bregma=0.5mm, lateral=1.1mm or bregma=-2.3mm, lateral=3mm). A collection catheter was then lowered stereotactically until cerebrospinal fluid was seen rushing into it to form a capillary head. This immediate feedback was possible thanks to the translucent nature of the PEEK tubing (Extruded Sub-Lite, Zeus) employed in the assembly of these microfabricated catheters. In the cases where this phenomenon was not visible, a second approach to the ventricle was attempted – either in the other hemisphere or in a different location within the same hemisphere. After validation of good catheter positioning, a three-component cement (Super-Bond C&B) was used to secure it in place together with a stainless steel headplate. The headplate was only added for head fixation in experiments in which the required sampling frequency was too high for repeated anaesthesia. Once the catheter was secured by the cement, it was removed from stereotactic support and a custom two-component protective chamber was placed around it. The base of the chamber was similarly cemented to the skull, whereas the cap was temporarily fixed by M1 screws. This design ensures the protection of the catheter and allows for multiple-animal housing. Before the cap is screwed on, the collection catheter was temporarily sealed by means of a custom filament plug and a two-component elastomer (Kwik-cast, World Precision Instruments). The weight of the protective chamber (∼0.6g) is low enough to not pose a burden for the animals and it is fabricated in a hard resin to prevent co-housed animals from damaging the implant.

#### Surgical recovery

All surgeries were performed under sterile conditions. Apart from the terminal procedures described above, all other surgical interventions required full recovery of the animals. To this end, Antisedan (50 µl, 5mg/mL) and Meloxicam (50 µl, Meloxidyl, 5 mg/mL) were administered intraperitoneally immediately after the surgery to ensure a fast recovery and to reduce pain and inflammation. The mice were then weighed and monitored for three days with a daily IP injection of Meloxicam.

### *In vivo* validation of the delivery micronozzle

#### Infusions of trypan blue dye

Wild-type mice were prepared for surgery as detailed above. A craniotomy was performed stereotactically corresponding to the coordinates of the target structure, either the AON (bregma=2.7mm, lateral=1mm, depth=2.6mm) or the striatum (bregma=-2.3mm, lateral=1.2mm, depth=4mm). One of three types of delivery catheters (i.e., polyimide FEC, micronozzle or pulled glass capillary) was then manually lowered to the desired location by means of a hydraulic actuator (Narishige). A 10-minute interval was implemented between the insertion of the catheter and the start of the infusion to allow for tissue relaxation. The infusion was controlled with a custom Python script which simultaneously commanded the syringe pump (33DDS - Harvard Apparatus), recorded data from the inline pressure sensor (uPS series – Labsmith) and displayed the progression in a graphic interface. A second 10-minute interval was then allowed between the end of the infusion and the retraction of the catheter. At the end of the procedure, each animal was euthanised by ketamine overdose and subsequently decapitated to allow for an immediate processing of the histological work.

#### Histology of trypan blue dye terminal infusions

In order to prevent the distribution of the dye to be significantly influenced by diffusive processes, the histological work was performed immediately after the termination of each infusion. Post-euthanasia, the non-perfused brain was extracted, and flash-frozen in dry ice (−80°C). It was then sectioned coronally at 100µm thickness with a cryostat (CM3050 S, Leica). All the slices containing visible dye were then mounted with a DAPI mounting medium and imaged with both bright-field and confocal microscopy.

#### Infusion cloud reconstruction

Comparison of anatomical features between the brain slices and the Allen atlas of the mouse brain was used to assign each slice a unique coordinate defining its position along the ventro-dorsal axis with respect to bregma. Each bright-field image was processed to remove any vignetting, it was white-balanced, straightened and cropped to the edges of the slice. Images were then binarized to obtain a logical map of the distribution cloud. This sequence of binary images was then assembled into a three-dimensional array after appropriate scaling based upon the coordinate of each slice. The resulting 3D volume was used to extract a surface for displaying purposes and to calculate the volume of distribution.

#### Target region coverage analysis

Independent of the injection method, desired brain coordinates are targeted with variable precision in different surgeries. To correct for this variability, we translated each infusion clouds to be centred in the centre of mass of the targeted brain area. The coverage of the target region was then evaluated according to three different metrics starting from the 3D volume of the target region and of the distribution cloud. The first metric, target coverage, indicates the proportion of the target region covered by the distribution cloud. The second, off-target ratio, indicates what proportion of the distribution cloud lies outside the target. Finally, the third metric, termed success of infusion (SOI) index, is a combination of the two previous values scaled between 0 and 1 according to the following equation:

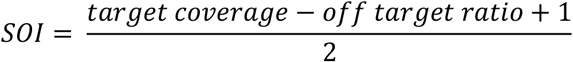

According to this definition, an SOI of 1 corresponds to two perfectly overlapping volumes whereas an SOI of 0 indicates two non-intersecting ones.

#### Olfactory exploration paradigm

The olfactory exploration paradigm was performed as described previously^18^. Briefly, we exposed head-restrained, male C57BL/6J mice (between 8-16 weeks old) to odorants that were either previously familiarized for 4 days or entirely novel during the session. Odorants were chosen from a stimulus set consisted of 20 diverse, single molecule substances (Sigma-Aldrich), commonly used in olfaction research (were anisole, ethyl valerate, eugenol, limonene, trymethyl pyrazine, benzyl acetate, cinnamaldehyde, pentenoic acid, linalyl formate, geraniol, thiophene, phenethyl alcohol, ethyl decanoate, nonadienal, acetic acid, benzaldehyde, heptanal, isoamyl acetate, octanoic acid, menthol).

During a session of the odorant exposure paradigm, a mouse experienced a randomized sequence of a total 4 familiar and 8 novel stimuli, each presented 10 times for 2 seconds. Between odorant stimulation, we introduced an intertrial interval drawn from an exponential distribution with an average of 30s and truncated at 2 minutes.

Respiration was monitored throughout using infrared thermography as described previously^20^. For analysis, breathing responses were normalized by subtracting the mean baseline breathing frequency prior to stimulus onset. Exploratory sniffing responses to the first three presentations of each odour were included in the analysis.

#### Glass capillary infusions

An injection of aCSF (150mM NaCl, 5mM KCl, 2mM NaH2PO4, 2.5mM CaCl2, 1mM MgCl2, 10mM HEPES, 10mM glucose, all from Sigma-Aldrich) was performed by a pulled glass capillary needle to deliver fluids to the AON. First, the mouse was head restrained in the setup, the silicone cap was removed from the skull and the craniotomy was opened again. Once the craniotomy was open, the cortex was moistened with sterile saline to prevent it from drying out. To track the injection path of the glass capillary needle, it was dipped in DiI (1,1’-Dioctadecyl-3,3,3’,3’-Tetramethylindocarbocyanine Perchlorate) prior to insertion into the brain. By the control of the micromanipulator, the glass capillary needle was then positioned just above the craniotomy and then lowered to a depth of 4 mm under bregma. After an interval of 10 minutes, the injection was started. Per hemisphere, 400-450nl were injected in batches of 50nl every 5 minutes. The total injection time was 45 minutes. All steps were performed under sterile conditions. After the injection and extraction of the glass capillary needle, the mouse performed the olfactory exploration paradigm.

#### Micronozzle infusions

Animals implanted with the microfabricated brain catheter were first placed in a head-restraining apparatus to allow for reliable access to the fluidic ports. The screw-on plugs were removed and an infusion line leading to a syringe pump (Harvard Apparatus 33dds) was connected. A low-volume bolus of infusate was delivered at minimally invasive flow rates (0.025µl/min) whilst the inline pressure was monitored via a microfluidic pressure sensor (uProcess uPS0250-C360-10 – LabSmith). This monitoring allows the interruption of the experiment in case the pressure exceeds a certain threshold value to guarantee the safeguard of the animal. Total infusion volumes were the same as in glass capillary injections. Upon the termination of the infusion, a 10-minute interval was employed to allow for the rebalancing of pressure within the infusion system. After this interval, the fitting was removed, the screw-on plug was replaced, and the mouse performed the olfactory exploration paradigm. Due to the presence of the chronic micronozzle implant, we were able perform an infusion with muscimol (Muscimol BODIPY TMR-X, red fluorescence) 2 days after aCSF injections to test the effect of silencing AON neural activity in our olfactory exploration paradigm. The procedure was as described above.

#### Histology

After mice performed their last session of the olfactory exploration paradigm, they were deeply anesthetized with ketamine-medetomidine, followed by transcardial perfusion with 0.1M phosphate-buffered saline (PBS) solution and 4% paraformaldehyde (PFA) solution. After 24 hours post-fixation in PFA, brains were stored in PBS for 2 days. Then they were sectioned in 60µm thick coronal slices with a vibratome (VT1000S, Leica). Free-floating sections were collected into 48 well plates filled with PBS. Aluminium foil was used to protect the slices from the light. Slides were mounted on glass microscopy slides with DAPI-containing mounting medium (4’,6-Diamidino-2-phenylindole dihydrochloride, Vectashield-H-1200-10, Sigma-Aldrich). All brain slices were imaged by a confocal microscope (LSM 710, Zeiss Group). By imaging the fluorescent muscimol and the DiI, the injection site could be visualized to determine if the muscimol injection was on target (AON, Franklin and Paxinos, 2008).

### *In vivo* validation of the sampling micronozzle

#### Sample collection

Sampling of cerebrospinal fluid requires immobilization of the animal. This was achieved either by head restraining (in animals where a head plate was implanted) or by gas anaesthesia (isoflurane). The catheter was uncovered by removing the two M1 screws holding the protective cap in place. After removal of the temporary rubber seal, CSF was sampled either with a micropipette or a pulled capillary and placed in an appropriate Eppendorf tube. At the end of the procedure, the catheter was resealed with the filament plug and the two-component rubber, and the protective cap was re-screwed in place. The volume of the liquid biopsy was estimated by the difference in weight between the empty Eppendorf (tare weight) and the filled one (gross weight) as measured by an analytical balance (CPA224S, Sartorius). The calculation was made under the assumption that, since CSF has a water content of ∼99%, its density can be approximated to 1mg/µl.

#### Glial fibrillary acid protein (GFAP) staining

Once CSF collection was no longer viable from one wild-type mouse at 74 weeks of age and 35 weeks post-implantation, it was euthanized and perfused. All procedures were conducted identically except for an extra two-step staining procedure performed in between the slicing and the mounting. For this procedure, the primary antibody was Gp-GFAP (173004, 1:1000, Synaptic Systems) and the secondary antibody was a Donkey anti-Guinea Pig Alexa 488 (706-545-148, 1:1000, Jackson Immunoresearch). All brain slices were then imaged with a confocal microscope (LSM 710, Zeiss Group) and the insertion track identified by visual inspection.

#### BBB integrity assessment with FITC-dextran

Thirty-five-week-old female C57BL/6J mice (n=3) were implanted with a CSF sampling catheter as described above. The coordinates of implantation chosen to grant access to the lateral ventricle were 0.7mm dorsally and 1mm laterally from bregma. All catheters were implanted at a depth of 2mm. One week post implantation the baseline barrier leakage was assessed by first injecting all three mice intravenously with 200µl of 25mg/ml 4kDa FITC-Dextran. The Dextran solution was prepared in aliquots in advance, stored at −20°C and shielded from light until minutes before the procedure. The animals were then placed under gas anaesthesia (isoflurane) so that they could be immobilized. Fifteen minutes after the IV injection a CSF sample was collected in an Eppendorf and subsequently stored in ice and away from light until further processing. Centrifugation (300 RCF) was then used to spin down the samples and 1µl of each was added to a separate well of a black 96-well plate together with 99µl of PBS. Fluorescence was then measured with a multi-mode microplate reader (FLUOstar^®^ Omega, BMG Labtech). The same pipeline was repeated one week later at two different timepoints (t=6 and t=24 hours) after the injection of LPS (1.5µg/µl per gram of bodyweight) to induce brain barrier leakage. All FITC-dextran injections after LPS induction used a different batch as the baseline measurement. One animal was found dead at the 24-hour timepoint and is therefore not reported in the analysis. Immediately after the 24-hour timepoint, mice were anaesthetised with a solution of a ketamine and xylazine and CSF isolation was also performed via cisterna magna extraction. Baseline CSF samples were also collected from the cisterna magna after FITC-dextran injection in two mice that received no catheter implant.

### Analysis of CSF biomarkers

#### Nucleic acids

A total of 18 CSF samples were analysed with a spectrophotometer (NanoDrop® ND-1000 UV-Vis Spectrophotometer) to measure nucleic acid concentrations. Of these 18 samples, 3 were negative controls (artificial CSF), 6 were obtained from one animal and 9 from a second one. The CSF samples from the first animal were collected over the span of four months and from the second over 6 months. In both cases, the minimum inter-sample interval was 24 hours. The ratio of sample absorbance at 260 and 280 nm for all samples was between 1.34 and 1.95 whereas the ratio at 260 and 230 nm ranged between 0.68 and 3.01.

#### Longitudinal sampling of inflammatory cytokines

One wild-type mouse was implanted with a CSF sampling catheter. CSF samples were collected on days 7, 16, 19, 21, 23 and 25 post-implantation following the procedure described above and kept frozen at −20°C. These liquid biopsies were then processed with a Cytokine Assay (Bio-Plex™) for the estimation of sample concentration of a total of 23 inflammatory cytokines (IL1-a, IL1-b, IL-2, IL-3, IL-4, IL-IL-6, IL-9, IL-10, IL-12p40, IL-12p70, IL-13, IL-14, Eotaxin, G-CSF, GM-CSF, IFN-g, KC, MCP-1, MIP1a, MIP1b, RANTES, TNF).

#### Longitudinal sampling of Aß40

Wild-type mice (n=3) between 13 and 22 weeks of age were implanted with CSF sampling catheters. Mice were housed on a 12h-12h light-dark cycle with food and water supplied ad libitum. CSF samples were collected either at the start (7 am) or at the end (7pm) of the light cycle with an inter-sample interval of 36 hours. Over 9 days, 3 samples per animal and per time of day (AM/PM) were collected for a total of 18 unique samples. After collection, the CSF biopsies were immediately frozen and kept at −-80°C until further processing. The volumes for all biopsies ranged between 4 and 15µl. Aβ1-40 levels in CSF were quantified by ELISA using Meso Scale Discovery (MSD, Meso Scale Diagnostics, Rockville, MD, USA) 96-well plates and in-house produced antibodies. MAb LTDA_Aβ40, which recognizes the C-terminus of Aβ1-40, was used as a capture antibody (coated ON, 4°C, 1,5µg/mL in PBS), and LTDA_rAβN (binding to the N-terminus of rodent Aβ) labelled with sulfoTAG as the detection antibody. Sample mix consisted of CSF dilutions in PBS 0.1% casein, mixed 1:1 with N2 rodent sulfo-TAG (1:2000 in PBS/casein), pre-incubated 15 min at RT, shaking 700 rpm, followed by ON incubation at 4°C on the coated MSD plates. Serial two-fold dilutions of rodent Aβ40, with concentrations ranging between 2.4 and 2500 pg/mL, were used as calibration curve. Sample concentrations were calculated based on the Aβ40 calibration curve using a non-linear fit, Log(agonist) vs response – variable slope (4 parameters, GraphPad Prism 9.3, GraphPad Software, San Diego, CA, USA). For the samples where the volume allowed (>10µl), up to two dilutions were made and the final concentration was taken as the mean. If the inferred concentration of a sample was below the sensitivity limit of the assay, the sample was excluded from the analysis. A normalisation was performed per animal by dividing by the mean of all samples.

## Acknowledgements

This work was supported by Interne Fondsen KU Leuven (C14/21/111 SH), VIB Grand Challenges Program “Targeting drugs to the brain” (SH), Fonds Wetenschappelijk Onderzoek-Vlaanderen (FWO-Flanders) doctoral fellowships 1S93620N (SM), 11K8122N (MLC) and grant G097022N (SH). Research in the Laboratory for Therapeutic and Diagnostic Antibodies was supported by Interne Fondsen KU Leuven.

