## Supplementary material for "Microfluidic interfaces for chronic bidirectional access to the brain"

### Appendix

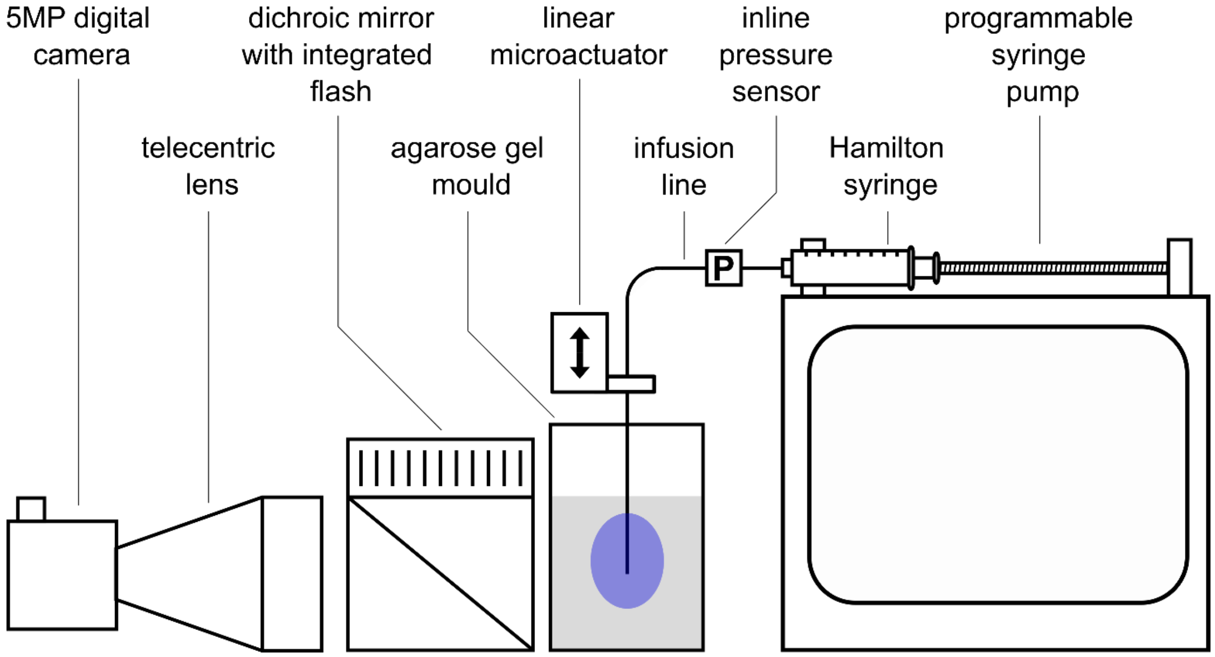


**Supplementary Figure 1. Setup for trypan blue dye infusions into agarose gel model.** A dichroic mirror is placed between the camera and the agarose gel mould to provide even lighting. The illumination is delivered by a flash and synchronised with the camera shutter. Catheters are placed in the mould with a linear microactuator at a constant speed of 4mm/s. A programmable syringe pump and an in-line pressure sensor are connected to a PC to respectively define the infusion protocol and monitor the progress of the infusion.

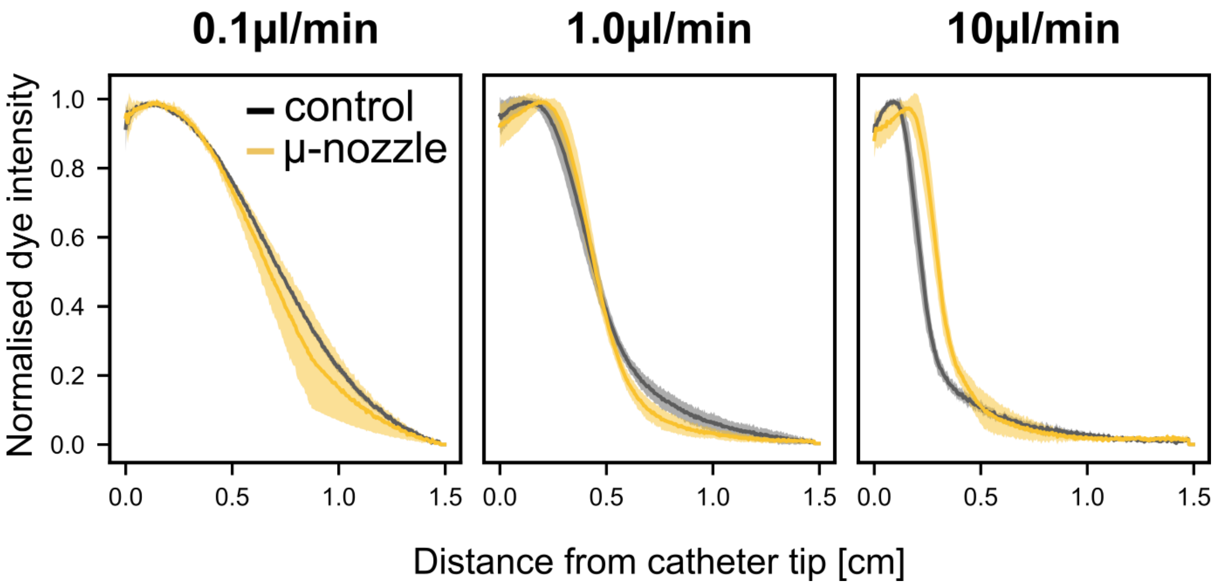


**Supplementary Figure 2. Dye intensity as function of the distance from the tip of the catheter.** Convective processes are predominant at higher flow rates, resulting in sharper edges of the distribution clouds. In very long infusions (such as the ones on the left), diffusion is an equally strong driver of the distribution of dye into agarose, resulting in very gentle slopes. A clear demarcation between infused and non-infused tissue is desirable in all applications where a leakage of the infusate beyond the target area can compromise the success of the procedure.

**Supplementary Table 1. Overview of protocols in low-volume delivery of trypan blue dye to agarose gel models.**

| Flow rate [µl/min] | Infusion volume [µl] | Repetitions |
|---|---|---|
| 0.1 | 0.5 | 2 |
| 0.1 | 2.0 | 2 |
| 0.2 | 2.0 | 2 |
| 0.5 | 2.5 | 2 |
| 1.0 | 5.0 | 2 |
| 2.0 | 5.0 | 2 |
| 5.0 | 5.0 | 2 |

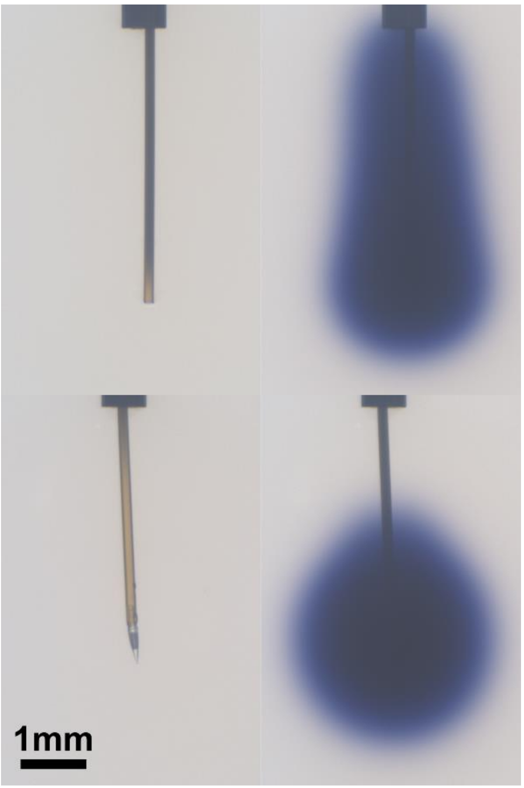


**Supplementary Figure 3. Example of low volume infusion of trypan blue dye in agarose gel brain model.** The start of the infusion is pictured on the left for both the EPC (control, top) and the micronozzle (bottom) whereas the respective ends are shown on the right. Both infusions pictured have the same protocol, a volume of infusion of 5 µl at a flow rate of 2 µl/min.

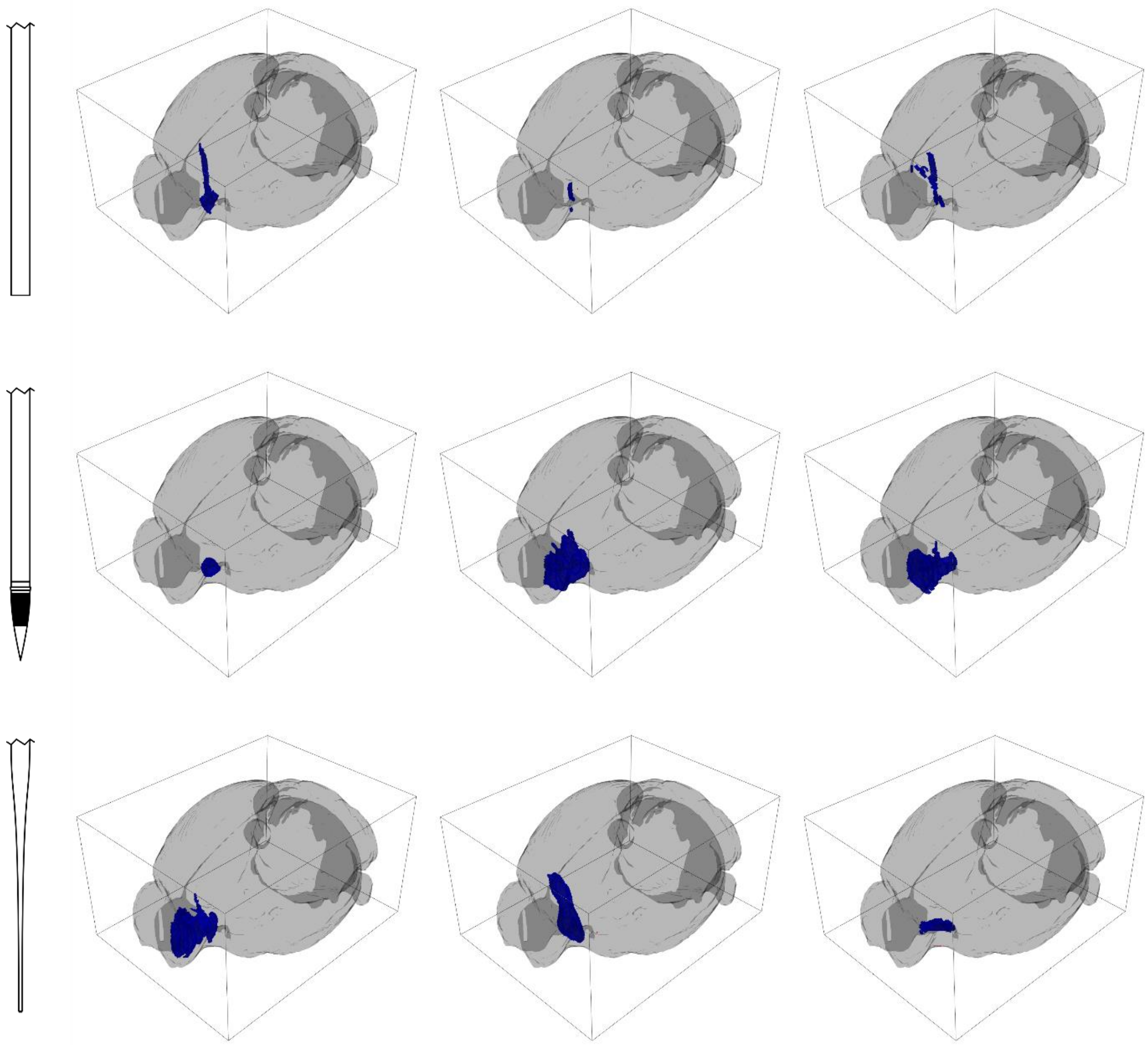

**Supplementary Figure 4. Overview of all infusions of Trypan blue in mice targeted to the Anterior Olfactory Nucleus.** Each row represents infusions performed with a different delivery system, EPC [top], micronozzle [middle] and pulled glass capillary [bottom]. Infusion volume was 400-450nl.

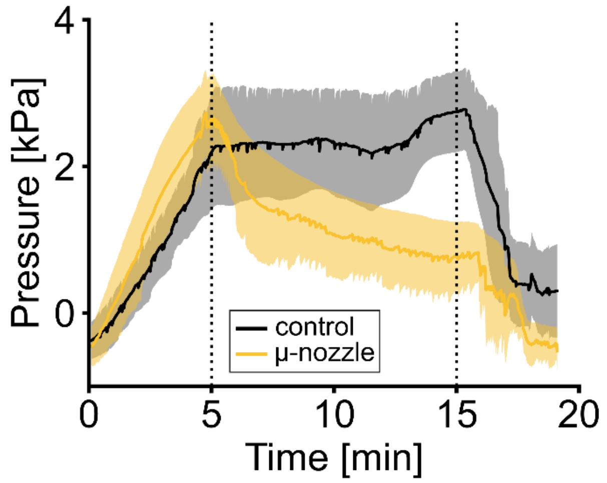


**Supplementary Figure 5. Pressure traces for the *in vivo* deliveries to the AON.** Data from the inline pressure sensor for *in vivo* infusions to the murine brain. Solid lines show the mean while shaded areas represent the standard error of the mean. The infusions are aligned and start at time t=0s. Dotted lines indicate, respectively, the end of the infusion and the retraction of the catheter. The release of pressure seen between infusion start and catheter retraction for the micronozzle can be attributed to a continuous release of fluid which is not seen in the control.

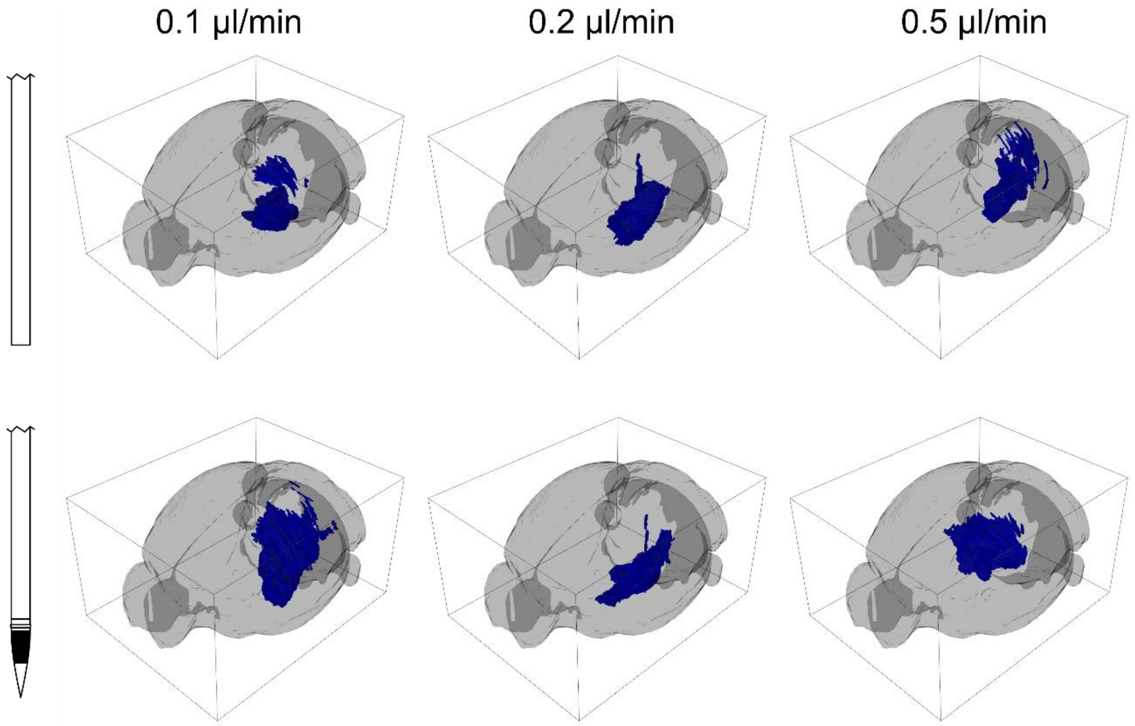


**Supplementary Figure 6. Overview of all trypan blue infusions in the striatum in mice.** Each row represents infusions with a different catheter system and each column represents a different flow rate. The volume of infusion in all cases was 2 µl.
